# Senescent Subtypes Transition Model

**DOI:** 10.1101/2023.06.10.544466

**Authors:** Ghanendra Singh

**Affiliations:** iCurious.in

## Abstract

Senescence has both beneficial and detrimental roles across the tissues over time. This dual nature is mediated by the senescence-associated secretory phenotype (SASP). Still, transient and persistent SASP secretion is poorly understood. There are some unknown mechanisms through which phenotypic transition takes place from beneficial helper (H) senescent cells to deleterious (D) senescent cell states. NOTCH1 is suggested to mediate switching between the different SASP secretomes. NOTCH1 also suppresses a transcription factor C/EBP*β* during the SASP secretion. Therefore, a hypothesis is proposed about the existence of negative feedback from C/EBP*β* to NOTCH1 together forming a senescence-switching circuit that might be mediating this phenotypic transition at the molecular level. Using a population dynamics model with competitive interaction to decipher the underlying transition mechanism between the senescent cell subtypes and a mechanistic model to explain the underlying molecular mechanisms of NOTCH1 signaling in modulating the two waves of SASP secretion. By designing effective senescence therapies with selective removal of deleterious senescent cells and maintaining sufficient helper senescent cells can enhance health span and reduce age-related effects.

## Introduction

Cellular senescence is an irreversible cell cycle arrest induced in response to stress [1]. Identified as a replication limit for the somatic cells after certain number of divisions [2]. Induction of senescence can have both beneficial or detrimental consequences depending on the contexts [3], nature such as oncogene [4], therapy [5], or replicative [2] induced, depending on the senescence phenotype and the dynamics of senescent cell clearance.

On the beneficial side, it promotes wound healing [6], tissue repair [7], tumor suppression [8], embryonic and fetal development [9]. Senescent cells may also play a role in the regeneration process [10]. A beneficial role for cellular senescence as an important modulator of dedifferentiation, a key mechanism for regeneration of complex structures. [11]. On the detrimental side, cellular senescence can also drive tumour growth and age-related disease progression [12].

### Mechanisms of SASP

An intracellular IL-1*α* miR-146*α*/*β* IL-6 C/EBP-*β* loop [13], contribute to the changes in gene expression that result in the SASP [14]. Positive feedback loops that reinforce the SASP and other regulatory mechanisms may contribute to enhanced SASP factor secretion. An example of such a self-amplifying positive feedback loop involves regulation of IL-1*α* secretion [15]. IL-1*α* is an upstream regulator of the IL-6/IL-8 cytokine network. IL-1*α* activates the transcription factors, C/EBP*β* and NF-*κ*, leading to enhanced production of IL-6 and IL-8 and IL-1*α* itself.

Time of induction from initiation to complete state of cellular senescence depends on cell type and the inducer [16]. SASP induction involves recruitment of BRD4 to senescence-activated super-enhancers adjacent to SASP factor genes [17]. Keratinocytes transiently exposed to SASP factors can have increased regenerative capacity. However, sustained exposure to the SASP can lead to the induction of senescence in these cells [18].

SASP can be tumor suppressive with induction of TGF-*β* or reinforcement with (IL6, IL8). SASP has a double edged sword in tumourigenesis [19]. Recently a stochastic simulation of senescence spread [20] identifies the differences in the number of signalling molecules secreted between subtypes of senescent cells can limit the spread of senescence.

SASP plays both pro and anti tumor roles. The pro-tumor and anti-tumor aspects of senescence may be reversible. Dormant cancer cells are quiescent but share some characteristics with a senescent cell. Likewise, quiescent and senescent cells share certain characteristics, and these states may exist on a spectrum [21]. TGF-*β* in the microenvironment induces a physiologically occurring immune suppressive senescent state [22]. It induces irreversible deep senescence with a 14-gene SASP. Deep senescence is a naturally occurring immune-suppressive cell state in cancer. NSCLC patients with high 14-gene SASP exhibit poor clinical outcome after ICI therapy. Identifying the molecular mechanisms and features of early vs. deep senescence across various cell types is key for potential beneficial vs. detrimental effects on health-span. Can we try to identify an adaptive mechanism for senolytic therapy delivery across heteregenous senescent cells?

### Senescence Subtypes

Senescence is a dynamic multistep process that is impacted by intra and extra cellular signals leading to heterogeneity in functional outcomes [23]. The anti-apoptotic mechanisms of senescent cells and SASP profiles are shaped continuously by the microenvironment, immune cells, and neighboring senescent cells [24]. Transient SASP secreting senescent cells are removed by the immune cells whereas persistent SASP secreting senescence cells have a pro-inflammatory, pro-apoptotic SASP inhibits the immune system[25].

Both the cell autonomous and non-autonomous properties of senescent cells are highly dependent on age and time. What determines the phenotype of senescent cells, and whether their effects on the tissue microenvironment are positive or negative? [26] Another interesting hypothesis proposed that senescent cells can be of helper (H) or deleterious (D) subtypes. These helper senescent cells may be convertible into deleterious senescent cells through an unknown mechanisms. Lowering the burden of these deleterious subtype below a specific threshold and minimise targeting the helper subtype to reduce its adverse effects may enhance health-span [27].

Deleterious senescent cells have a pro-inflammatory, pro-apoptotic, tissue-damaging SASP and are susceptible to senolytics. Helper senescent cells do not have a SASP and are not cleared by senolytics. Helper senescent cells appear to promote stem and progenitor cell determination into appropriate lineages and functions and have a beneficial impact on tissue homoeostasis. Its important to understand the heterogeneity of senescence [28]. How does these helper and deleterious senescent subtype population cells interact with each other?

### Mechanistic Switching

While the beneficial versus detrimental implications of the senescence-associated secretome remain an issue of debate, time-resolved analyses of its composition, regulatory mechanisms and functional consequences have been largely missing. The dynamic activity of NOTCH1 is now shown to direct two distinct senescence phenotypes, by first promoting a pro-senescent TGF-*β* dependent secretome, followed by a second wave of pro-inflammatory, senescence-clearing cytokines. NOTCH is a spatiotemporal regulator of SASP composition in senescence cells [29].

NOTCH1 mediated secretome switching suggests that the dynamic nature of NOTCH1 activity contributes to transition between these two SASPs during senescence [30]. NOTCH1 signaling reciprocally regulates inflammatory cytokines and TGF-*β* during senescence. During primary senescence, NOTCH1 drives TGF-*β* while simultaneously suppressing the pro-inflammatory SASP by repressing C/EBP*β*. The NOTCH1 signaling inhibits IL-1*α* by down regulating C/EBP*β*. At secondary senescence, NOTCH activity is down-regulated and C/EBP*β* is upregulated allowing expression of the pro-inflammatory SASP. NOTCH1 also regulate chromatin structure during senescence [31]. How NOTCH1 activity is downregulated at later stages of senescence, despite the high levels of NOTCH1 remains unclear. Does NOTCH1 mediated switch between two different SASPs have any significant functional role?

## Results

### Population dynamics of helper and deleterious senescent cells

Senescent cells may be of helper (H) or deleterious (D) subtypes depending on the context and type of inducer of cellular senescence. It is critically important to understand the diverse populations of senescent cells and their functional relevance across tissues and diseases. Important to identify strategies for selectively removing deleterious senescent cells at the right time points, while leaving helper senescent cells available for physiological functions such as tissue repair.

Helper senescent cells might transition into deleterious subtype through as yet unknown mechanisms that need to be explored [27]. Understanding if local or systemic clearance of senescent cells may be the better option for aged or diseased patients. Complete removal of helper senescent cell can impair physiological functions.

Selectively lower the burden of persistent, pro-inflammatory, pro-apoptotic deleterious senescent cells, while retaining beneficial helper senescent cells, or at least preserving ability for helper cells to be generated when needed is key to tackle age related diseases. Therefore, proposing a Lotka Volterra based population dynamics model [32] for senescent cell subtypes with phenotypic transition, competition, and senolytic therapy for selective removal of deleterious senescent cells. Assumption is that the competitive effect existing between H and D cells along with secretion of SASP in transient and persistent manner and their effect on senescent subtypes. Below, system of ODEs representing population dynamics of senescent cell subtypes with possible mechanism of inter convertibility from H to D subtypes as follows.

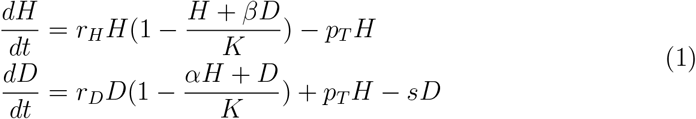

In the above equation 1, H represents the helper senescent cell population, D represents the deleterious senescent cell population, *r*_*H*_ is the helper cells accumulation rate, *r*_*D*_ is the deleterious cells accumulation rate, K is carrying capacity within appropriate tissue, *α* and *β* are competition parameters between helper and deleterious population subtypes, *p*_*T*_ is phenotypic transition rate from helper to deleterious subtype and *s* is the external senolytic dosage applied on deleterious subtype cells.

Figure 1a shows a population dynamics diagram of helper (H) and deleterious (D) senescent cells mutually inhibiting each other and transition from D to H during persistent SASP. Modeling dynamics shown in the figure 1b matches with the dual SASP waves described in [33]. Senescence induction during the first wave initially indicates the role of beneficial helper senescence cells due to the transient SASP nature. But later due to persisitant SASP over the time, deleterious senescence cells cross a particular threshold due to phenotypic transition and takes over the beneficial senescent cell population during the second wave [30].

**Figure 1:**
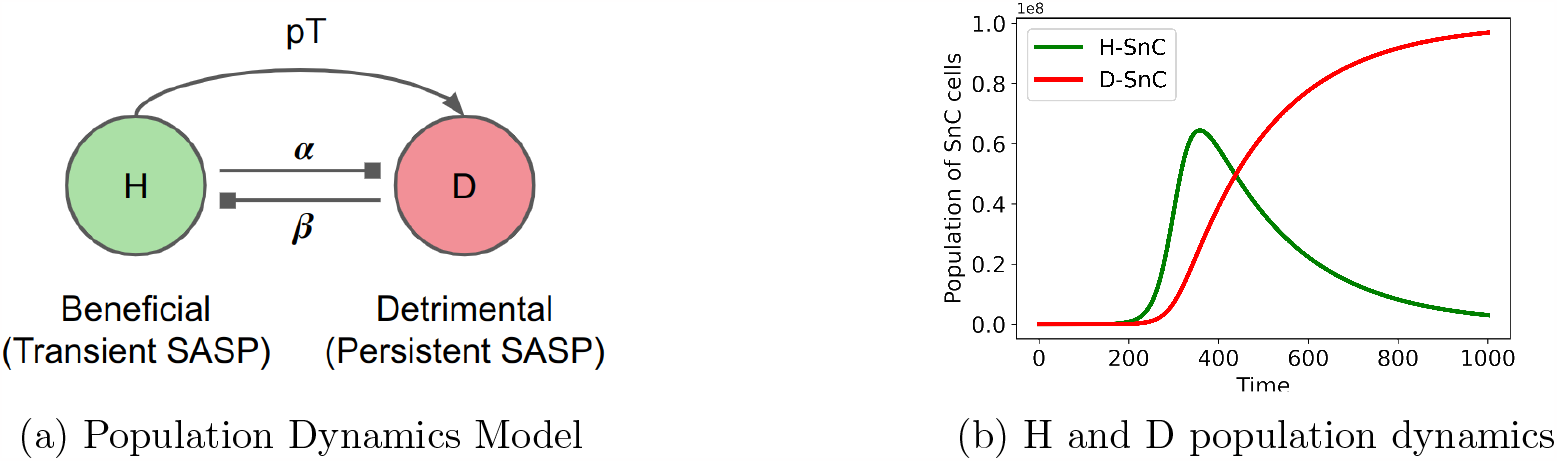
Population dynamics of helper and deleterious senescent cells. (a) H and D senescent cell population dynamics model with phenotypic transition *p*_*T*_ and competition parameters *α* and *β*. (b) Temporal dynamics of the model. Initially H senescent cells exist due to transient SASP with beneficial nature. Over time, D senescent cells takes over due to persistent SASP with detrimental nature. Parameters of the model are available in the Supplementary file.

Initially helper senescent subtypes exists and over the time due to persistent SASP feedback loop nature, possibly undergoes phenotypic transition into deleterious senescent subtypes. Figure 2a shows phenotypic transition *p*_*T*_ from helper (H) into deleterious (D) senescent cell population under normal condition. When *p*_*T*_ rate is low in nature, helper (H) senescent cells population persists for longer duration till the onset of D population as shown in figure 2b. Similarly, during high *p*_*T*_ rates, D population quickly takes over H population type leading to early onset of detrimental effects of senescence as shown in figure 2c.

**Figure 2:**
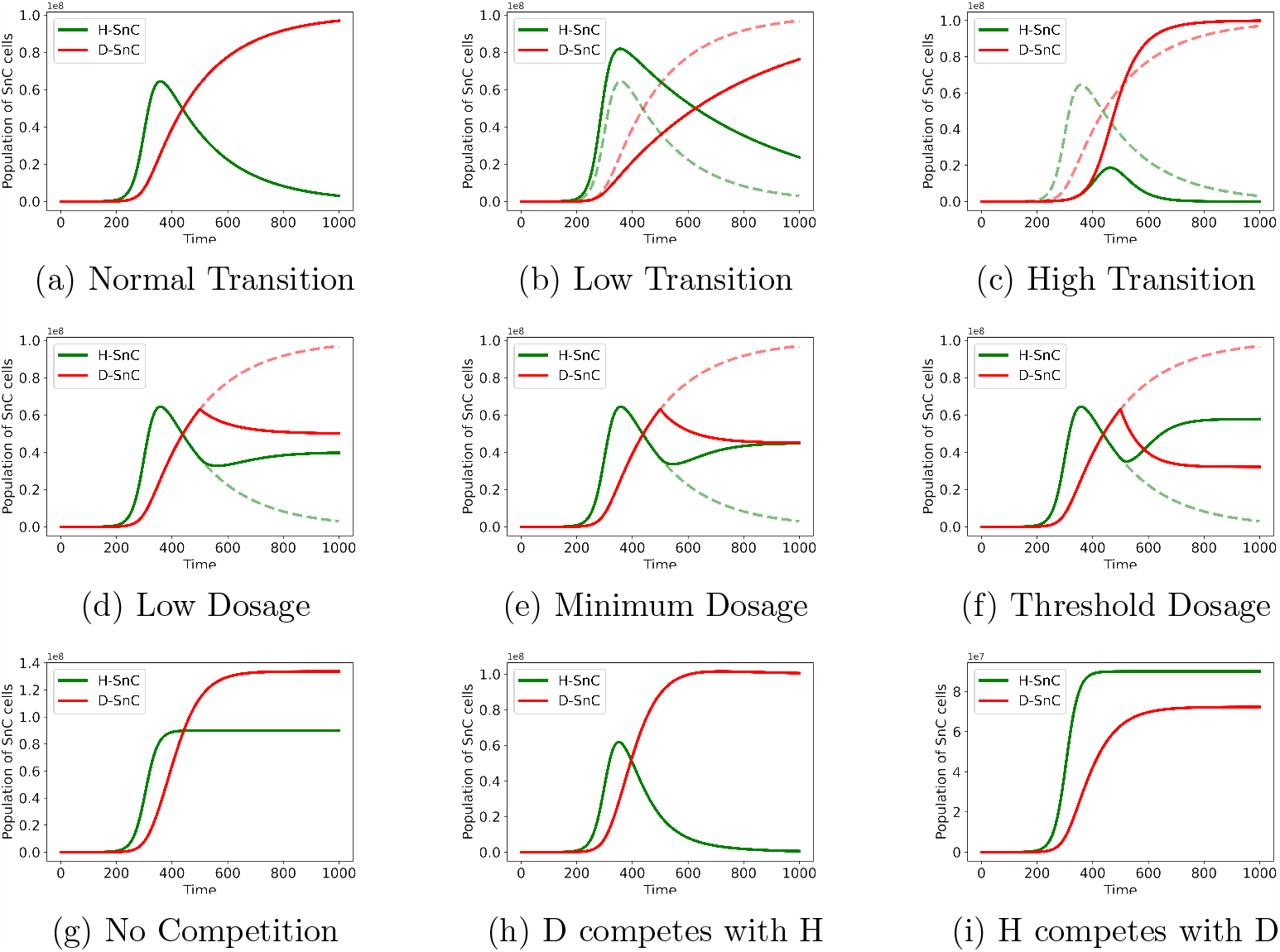
Population dynamics of Helper (H) and Deleterious (D) Senescent Cells with Phenotypic Transition, Senolytic Dosage, and Competition among them. Senescence cells undergoes phenotypic transition (PT) from H to D subtypes at normal rate(*pt*=0.005) in (a), low rate(*pt*=0.002) in (b), and high rate(*pt*=0.02) in (c). Dynamics on application of senolytic for selective clearance of D cells for low dosage (d), minimum dosage (e), and threshold dosage (f). No competition between H and D cells (g), D competes with H cells (h), and H competes with D cells (i).

**Figure 3:**
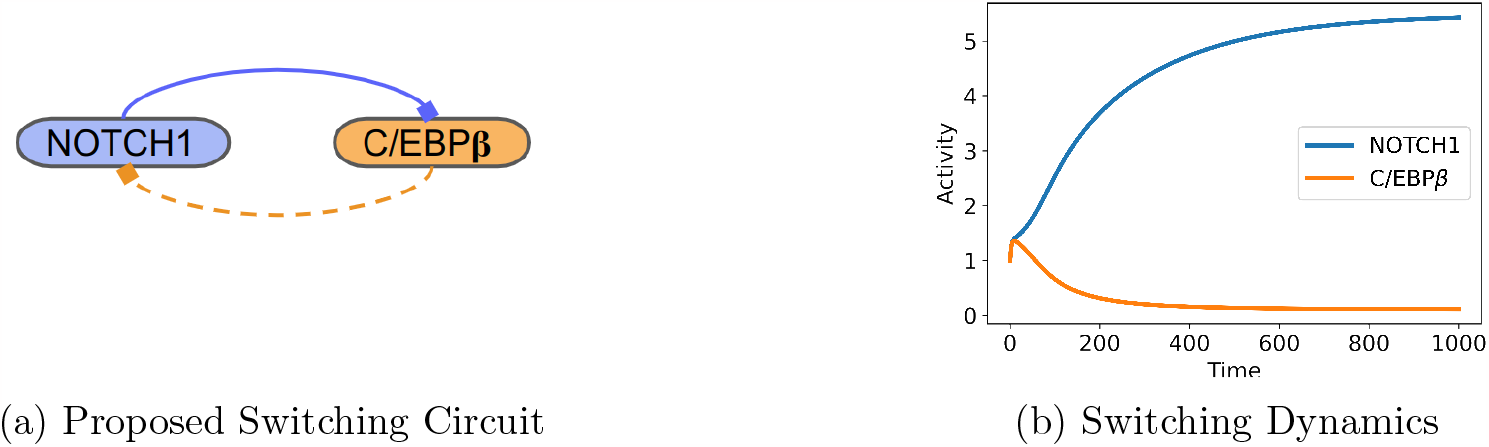
Hypothesized NOTCH1-C/EBP*β* switching circuit and its dynamics. (a) Proposed negative feedback from C/EBP*β* to NOTCH1 together forming a switching circuit. (b) Switching Dynamics of the circuit. NOTCH1 is

**Figure 4:**
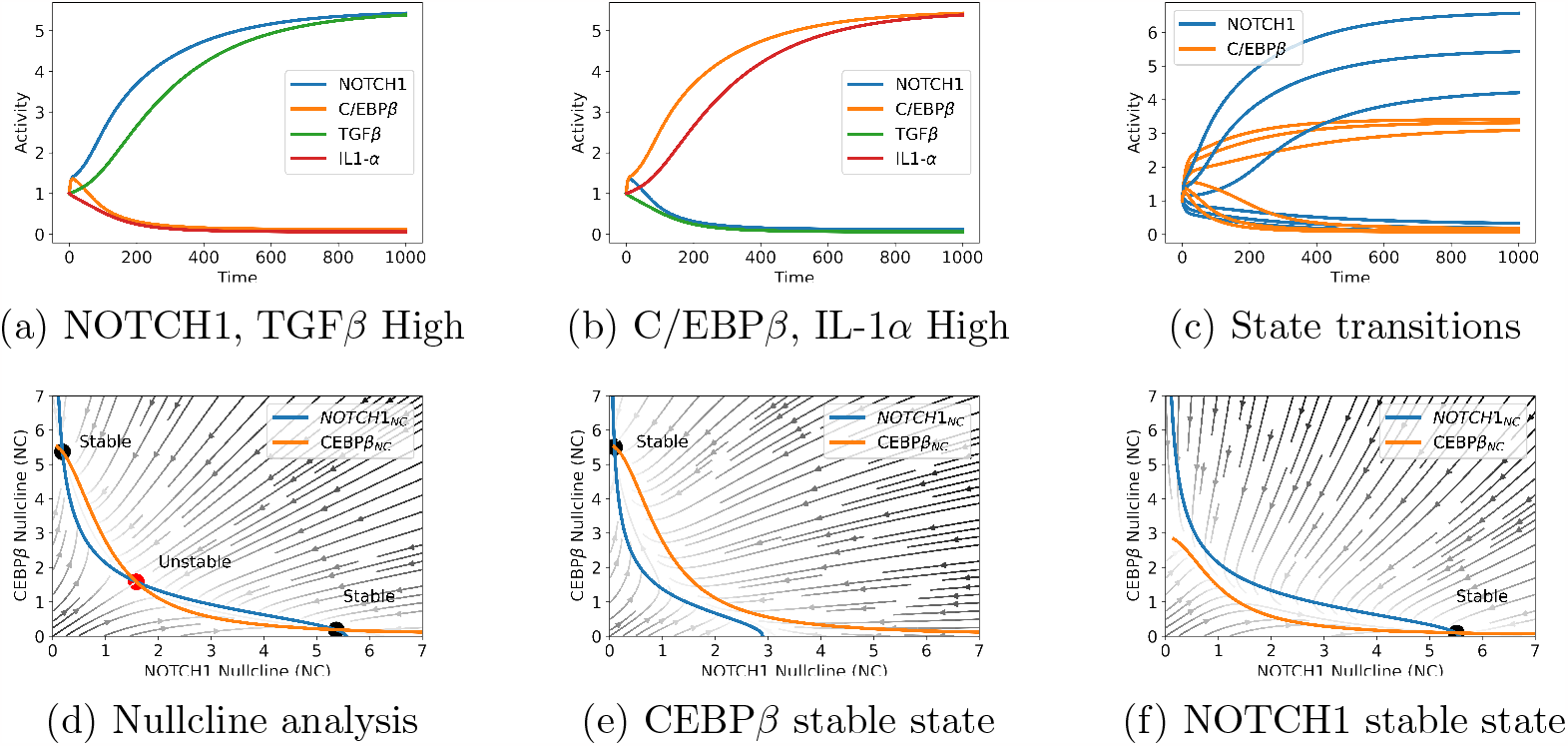
Nullcline and bistability dynamics of NOTCH1 and C/EBP*β* switching. (a) NOTCH1, TGF*β* are high and C/EBP*β*, IL-1*α* are low (*d*_*T*_ =0.01 and *d*_*I*_=0.02). (b) C/EBP*β*, IL-1*α* are high and NOTCH1, TGF*β* are low (*d*_*T*_ =0.02 and *d*_*I*_=0.01). (c) Varying *β*_1_ from 0.1 to 0.7 shows transition for both NOTCH1 and C/EBP*β*. Nullcline analysis is shown in the next three figures. (d) Two stable states, one each for NOTCH1 and C/EBP*β* one unstable state might exist based on the proposed hypothesis. (e) Stable steady state exists for C/EBP*β* at (*d*_*T*_ =0.04 and *d*_*I*_=0.01). Similarly, (f) stable steady state exists for NOTCH1 at (*d*_*T*_ =0.01 and *d*_*I*_=0.01). Params: *β*_*N*_ =0.5, *β*_*C*_=0.5, n=2, *d*_*N*_ =0.2, *d*_*C*_=0.2, *p*_*T*_ =0.01, *p*_*I*_=0.01, *d*_*T*_ =0.01, *d*_*I*_=0.01, *r*_*T*_ =0.11, *r*_*I*_=0.11, and all the initial value for four (N,T,C,I) were set to 1. Parameters are based on trial and error to build this model.

**Figure 5:**
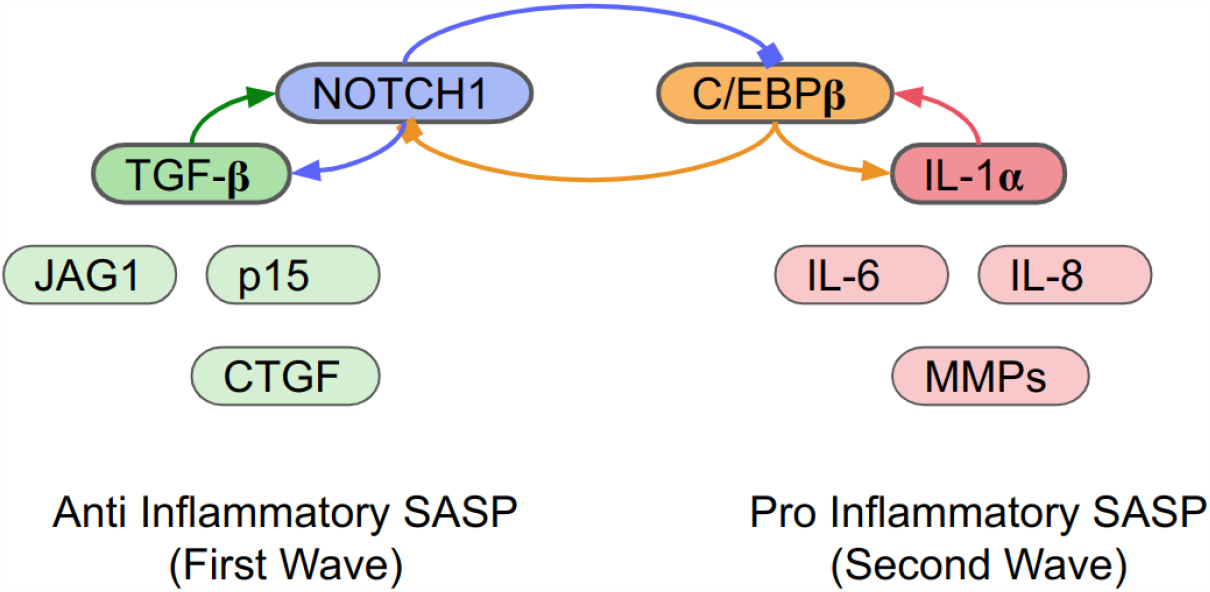
NOTCH1 C/EBP*β* Switching circuit with positive feedback loops. Two state switching model of the SASP. Existence of a toggle switch directly between NOTCH1 and CEBP*β* or through other unknown transcription factor downstream of CEBP*β* possibly. A positive feedback loops is established through TGF*β* and IL-1*α*.

For effective senolytic dosage for selective removal of deleterious (D) subtype from the senescent cell population, consider a simple factor (s) representing the senolytic based removal of D subtypes. For low senolytic dosage, D subpopulation is higher than H subtypes as shown in the figure 2d. In figure 2e, D is nearly equal to the H for minimum senolytic dosage and H subtype population is higher than D subtype population using effective threshold senolytic dosage (TSD) in the figure 2f.

Local senolysis can only partially replicates the benefits seen at the systemic senolysis [34]. Senescent cells through their SASP, lead to senescence in distant cells. It indicates that for optimizing senolytic drugs, it may require systemic instead of local senescence cells clearance targeting to extend healthy aging. Therefore, it might be beneficial if deleterious senescence cells and its effects can be reduced or regulated at the systems level. It requires further confirmatory experimental investigations.

Considering competition between the helper (H) and deleterious (D) subtype senescent cell population. When no competition exists (*α*=0 and *β*=0), both the sub populations reaches a steady state with D higher than H as shown in the figure 2g. In figure 2h, assuming, D senescent cell population competes with H cells, it might leads to D cells taking over H cells after a certain time frame. Lastly, if we assume that H competes with the D senescent cell population, both subtype population exists with H higher than D subtypes as shown in the figure 2i which is highly impossible based on the experimental evidences.

### Molecular mechanisms of Senescent Subtype switching

Senescent cells accumulate over time and can have both beneficial and detrimental effects through the secretion of factors called as the senescence-associated secretory phenotype (SASP) [35] or senescence-messaging secretome [36]. These two different effects are linked to paracrine signaling through the SASP in which senescent cells produce a collection of factors including cytokines, growth factors and matrix-modifying enzymes, NF-kB and C/EBPb [37]. From beneficial properties perspective, SASP factors recruit natural killer cells to eliminate malignant cells [38]. Senescent cells also recruit innate immune cells to kill tumour cells [39, 40]. Based on phenotypic states, two states has been hypothesized to exist in non-immune senescent cells [16]. Immunogenic senescent cells release immune cell-attracting SASP factors and express surface markers that promote clearance of the senescence cells. Senescence cells in an “anergic state” do not stimulate immune responses [41].

Composition of the SASP is dynamically and spatially regulated. Dynamic variation of NOTCH1 activity during senescence indicates a functional balance between these two distinct secretomes namely transient and persistent SASP. Secretome is spatiotemporally regulated by the NOTCH1 [30]. Changing composition of the SASP can determine the beneficial and detrimental aspects of the senescence program, shifting the balance to either an immunosuppressive/profibrotic environment or proinflammatory/fibrolytic state [29]. Both NOTCH1 and TGF*β* forms a regulatory positive feedback loop that inhibits the activity of prostate basal stem dormancy [42]. Senescence induced by the oncogene RAS, NOTCH1 signaling orchestrated a pre-SASP secretome. Initially, NOTCH1 is high and suppresses C/EBP*β* IL-1*α* leading to the SASP which does not attract immune cells. Over the time, NOTCH1 activity is suppressed, giving rise to C/EBP*β* IL-1*α* and classical SASP inflammatory cytokines [43]. Secondary senescence phenotype of the senescent cells is also mediated by NOTCH1 [44]. Also, time is a key player for these distinct SASP with beneficial and detrimental effects [45].

Hence, hypothesizing about the existence of a double negative feedback loop or toggle switching behavior between NOTCH-1 and C/EBP*β* directly or indirectly through intermediate transcription factor which might enable expression of TGF-*β* and IL-1*α* in an alternate manner and vice versa. Therefore, proposing a mechanistic model underlying senescence phenotypic switching between helper and deleterious senescent cell subtypes. System of equations of the proposed model is given below:

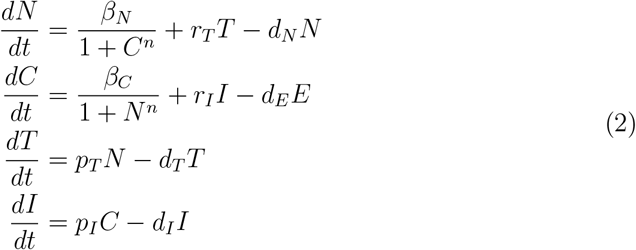

In equation 2, N is NOTCH1, C is C/EBP*β*, T is TGF*β*, and I is IL-1*α*. For NOTCH1: *β*_*N*_ production rate, *d*_*N*_ degradation rate, and *r*_*T*_ TGF-*β* induced NOTCH1 production rate. For C/EBP*β*: *β*_*C*_ production rate, *d*_*C*_ degradation rate, *r*_*I*_ IL-1*α* induced production rate. Production *p*_*T*_ and degradation rate *d*_*T*_ for TGF-*β*. Production rate *p*_*I*_ and degradation rate *d*_*I*_ for IL-1*α*.

## Conclusions

Paper focuses on the beneficial and detrimental role of senescent cells to highlight the plausible underlying conditions under which the senescence dual behavior comes into the picture. Hypothesized about the existence of a negative feedback from C/EBP*β* to NOTCH1 using a mechanistic model and a population dynamics model to decipher the underlying transition mechanism of helper senescence cell phenotype into detrimental senescence cells. Proposed switching circuit and hypothesis needs to be verified experimentally.

In essence, how does senescence subtype phenotypic switching takes place from helper to detrimental senescent cells is discussed at the population level and also at the molecular level using mathematical modeling. It highlights the variations seen in senescence subtype transition rates might determine the nature of transient or persistent SASP for beneficial or detrimental effects of senescence. How effective dosage and competition techniques might help in regulating deleterious effects and promoting beneficial effects of senescence? Next from mechanistic perspective, possibly the first SASP wave initially exist due to the abundance of helper senescent cell population. Then, second SASP wave may occurs when the deleterious senescent cells population crosses a certain threshold due to phenotypic transition and takes over the helper population. The dynamic activity of NOTCH is shown to direct two distinct senescence phenotypes, by first promoting a pro-senescent TGF-*β*-dependent secretome, followed by a second wave of pro-inflammatory, senescence-clearing cytokines. Finally, tried to map the simulation results with the experimental finding shown about the role of NOTCH1 in mediating a switch between two distinct secretomes during senescence [30]. By understanding the context dependent effects of cellular senescence, it might help in limiting the detrimental effects while maintaining its beneficial effects, and develop novel therapeutics to prevent age related diseases [3].

Lastly, double negative feedback loop might be the possible mechanism behind NOTCH1 mediated regulation of TGF*β* and C/EBP*β* during senescence and its disruption may contribute to aging related diseases as asked by [29]. Also, it would be interesting to see how dynamic, time dependent transient or persistent SASP regulates the spread of senescence [20]. Does NOTCH1 have any significant role in the difference seen in secreted signaling molecules within the senescence subtypes depending upon transition mechanisms which might be limiting the growth of specific or overall senescence cell population.

## Acknowledgement

I would like to thanks all the researchers on whose shoulders this work was carried out to develop a phenomenological senescence subtype switching model. Research is part of the iCurious.in organisation to contribute in open ageing research.

## References

[1] J. Campisi, “Aging, cellular senescence, and cancer,” Annual review of physiology, vol. 75, pp. 685–705, 2013.

[2] L. Hayflick, “The limited in vitro lifetime of human diploid cell strains,” Experimental cell research, vol. 37, no. 3, pp. 614–636, 1965.

[3] P. Lecot, F. Alimirah, P.-Y. Desprez, J. Campisi, and C. Wiley, “Contextdependent effects of cellular senescence in cancer development,” British journal of cancer, vol. 114, no. 11, pp. 1180–1184, 2016.

[4] C. Michaloglou, L. C. Vredeveld, M. S. Soengas, C. Denoyelle, T. Kuilman, C. M. Van Der Horst, D. M. Majoor, J. W. Shay, W. J. Mooi, and D. S. Peeper, “Brafe600-associated senescence-like cell cycle arrest of human naevi,” Nature, vol. 436, no. 7051, pp. 720–724, 2005.

[5] J. A. Ewald, J. A. Desotelle, G. Wilding, and D. F. Jarrard, “Therapy-induced senescence in cancer,” JNCI: Journal of the National Cancer Institute, vol. 102, no. 20, pp. 1536–1546, 2010.

[6] M. Demaria, N. Ohtani, S. A. Youssef, F. Rodier, W. Toussaint, J. R. Mitchell, R.-M. Laberge, J. Vijg, H. Van Steeg, M. E. Dollé, et al., “An essential role for senescent cells in optimal wound healing through secretion of pdgf-aa,” Developmental cell, vol. 31, no. 6, pp. 722–733, 2014.

[7] K.-H. Kim, C.-C. Chen, R. I. Monzon, and L. F. Lau, “Matricellular protein ccn1 promotes regression of liver fibrosis through induction of cellular senescence in hepatic myofibroblasts,” Molecular and cellular biology, vol. 33, no. 10, pp. 2078–2090, 2013.

[8] G. P. Dimri, K. Itahana, M. Acosta, and J. Campisi, “Regulation of a senescence checkpoint response by the e2f1 transcription factor and p14arf tumor suppressor,” Molecular and cellular biology, vol. 20, no. 1, pp. 273–285, 2000.D.

[9] Muñoz-Espín, M. Cañamero, A. Maraver, G. Gómez-López, J. Contreras, S. Murillo-Cuesta, A. Rodríguez-Baeza, I. Varela-Nieto, J. Ruberte, M. Collado, et al., “Programmed cell senescence during mammalian embryonic development,” Cell, vol. 155, no. 5, pp. 1104–1118, 2013.

[10] M. H. Yun, H. Davaapil, and J. P. Brockes, “Recurrent turnover of senescent cells during regeneration of a complex structure,” elife, vol. 4, p. e05505, 2015.

[11] H. E. Walters, K. E. Troyanovskiy, A. M. Graf, and M. H. Yun, “Senescent cells enhance newt limb regeneration by promoting muscle dedifferentiation,” Aging Cell, p. e13826, 2022.

[12] D. J. Baker, B. G. Childs, M. Durik, M. E. Wijers, C. J. Sieben, J. Zhong, R. A. Saltness, K. B. Jeganathan, G. C. Verzosa, A. Pezeshki, et al., “Naturally occurring p16ink4a-positive cells shorten healthy lifespan,” Nature, vol. 530, no. 7589, p. 184–189, 2016.

[13] D. Bhaumik, G. K. Scott, S. Schokrpur, C. K. Patil, A. V. Orjalo, F. Rodier, G. J. Lithgow, and J. Campisi, “Micrornas mir-146a/b negatively modulate the senescence-associated inflammatory mediators il-6 and il-8,” Aging, vol. 1, no. 4, p. 402, 2009.

[14] T. Tchkonia, Y. Zhu, J. Van Deursen, J. Campisi, J. L. Kirkland, et al., “Cellular senescence and the senescent secretory phenotype: therapeutic opportunities,” The Journal of clinical investigation, vol. 123, no. 3, p. 966–972, 2013.

[15] A. V. Orjalo, D. Bhaumik, B. K. Gengler, G. K. Scott, and J. Campisi, “Cell surface-bound il-1α is an upstream regulator of the senescence-associated il-6/il-8 cytokine network,” Proceedings of the National Academy of Sciences, vol. 106, no. 40, p. 17031–17036, 2009.

[16] A. Hernandez-Segura, T. V. de Jong, S. Melov, V. Guryev, J. Campisi, and M. Demaria, “Unmasking transcriptional heterogeneity in senescent cells,” Current Biology, vol. 27, no. 17, p. 2652–2660, 2017.

[17] N. Tasdemir, A. Banito, J.-S. Roe, D. Alonso-Curbelo, M. Camiolo, D. F. Tschaharganeh, C.-H. Huang, O. Aksoy, J. E. Bolden, C.-C. Chen, et al., “Brd4 connects enhancer remodeling to senescence immune surveillancebrd4 controls senescence immune surveillance,” Cancer discovery, vol. 6, no. 6, p. 612–629, 2016.

[18] B. Ritschka, M. Storer, A. Mas, F. Heinzmann, M. C. Ortells, J. P. Morton, O. J. Sansom, L. Zender, and W. M. Keyes, “The senescence-associated secretory phenotype induces cellular plasticity and tissue regeneration,” Genes & development, vol. 31, no. 2, p. 172–183, 2017.

[19] N. Frey, S. Venturelli, L. Zender, and M. Bitzer, “Cellular senescence in gastrointestinal diseases: from pathogenesis to therapeutics,” Nature Reviews Gastroenterology & Hepatology, vol. 15, no. 2, p. 81–95, 2018.

[20] L. K. Martin, L. Schumacher, and T. Chandra, “Modelling the dynamics of senescence spread,” bioRxiv, p. 2023–03, 2023.

[21] K. Truskowski, S. R. Amend, and K. J. Pienta, “Dormant cancer cells: programmed quiescence, senescence, or both?,” Cancer and Metastasis Reviews, p. 1–11, 2023.

[22] S. Matsuda, A. Revandkar, T. D. Dubash, A. Ravi, B. S. Wittner, M. Lin, R. Morris, R. Burr, H. Guo, K. Seeger, et al., “Tgf-β in the microenvironment induces a physiologically occurring immune-suppressive senescent state,” Cell Reports, vol. 42, no. 3, 2023.

[23] N. Herranz, J. Gil, et al., “Mechanisms and functions of cellular senescence,” The Journal of clinical investigation, vol. 128, no. 4, p. 1238–1246, 2018.

[24] L. G. L. Prata, I. G. Ovsyannikova, T. Tchkonia, and J. L. Kirkland, “Senescent cell clearance by the immune system: Emerging therapeutic opportunities,” in Seminars in immunology, vol. 40, p. 101275, Elsevier, 2018.

[25] J. Kirkland and T. Tchkonia, “Senolytic drugs: from discovery to translation,” Journal of internal medicine, vol. 288, no. 5, p. 518–536, 2020.

[26] M. Demaria, P. Y. Desprez, J. Campisi, and M. C. Velarde, “Cell autonomous and non-autonomous effects of senescent cells in the skin,” Journal of Investigative Dermatology, vol. 135, no. 7, p. 1722–1726, 2015.

[27] U. Tripathi, A. Misra, T. Tchkonia, and J. L. Kirkland, “Impact of senescent cell subtypes on tissue dysfunction and repair: importance and research questions,” Mechanisms of Ageing and Development, vol. 198, p. 111548, 2021.

[28] R. L. Cohn, N. S. Gasek, G. A. Kuchel, and M. Xu, “The heterogeneity of cellular senescence: Insights at the single-cell level,” Trends in cell biology, 2022.

[29] Y. Ito, M. Hoare, and M. Narita, “Spatial and temporal control of senescence,” Trends in Cell Biology, vol. 27, no. 11, p. 820–832, 2017.

[30] M. Hoare, Y. Ito, T.-W. Kang, M. P. Weekes, N. J. Matheson, D. A. Patten, S. Shetty, A. J. Parry, S. Menon, R. Salama, et al., “Notch1 mediates a switch between two distinct secretomes during senescence,” Nature cell biology, vol. 18, no. 9, p. 979–992, 2016.

[31] A. J. Parry, M. Hoare, D. Bihary, R. Hänsel-Hertsch, S. Smith, K. Tomimatsu, E. Mannion, A. Smith, P. D’Santos, I. A. Russell, et al., “Notch-mediated noncell autonomous regulation of chromatin structure during senescence,” Nature communications, vol. 9, no. 1, p. 1840, 2018.

[32] P. J. Wangersky, “Lotka-volterra population models,” Annual Review of Ecology and Systematics, vol. 9, p. 189–218, 1978.

[33] C. A. Schmitt, “The persistent dynamic secrets of senescence,” Nature Cell Biology, vol. 18, no. 9, p. 913–915, 2016.

[34] J. N. Farr, D. Saul, M. L. Doolittle, J. Kaur, J. L. Rowsey, S. J. Vos, M. N. Froemming, A. B. Lagnado, Y. Zhu, M. Weivoda, et al., “Local senolysis in aged mice only partially replicates the benefits of systemic senolysis,” The Journal of clinical investigation, vol. 133, no. 8, 2023.

[35] J.-P. Coppé, C. K. Patil, F. Rodier, Y. Sun, D. P. Muñoz, J. Goldstein, P. S. Nelson, P.-Y. Desprez, and J. Campisi, “Senescence-associated secretory phenotypes reveal cell-nonautonomous functions of oncogenic ras and the p53 tumor suppressor,” PLoS biology, vol. 6, no. 12, p. e301, 2008.

[36] T. Kuilman and D. S. Peeper, “Senescence-messaging secretome: Sms-ing cellular stress,” Nature reviews cancer, vol. 9, no. 2, p. 81–94, 2009.

[37] J.-P. Coppé, P.-Y. Desprez, A. Krtolica, and J. Campisi, “The senescence-associated secretory phenotype: the dark side of tumor suppression,” Annual review of pathology: mechanisms of disease, vol. 5, p. 99–118, 2010.

[38] A. Iannello, T. W. Thompson, M. Ardolino, S. W. Lowe, and D. H. Raulet, “p53-dependent chemokine production by senescent tumor cells supports nkg2ddependent tumor elimination by natural killer cells,” Journal of Experimental Medicine, vol. 210, no. 10, p. 2057–2069, 2013.

[39] A. Ventura, D. G. Kirsch, M. E. McLaughlin, D. A. Tuveson, J. Grimm, L. Lintault, J. Newman, E. E. Reczek, R. Weissleder, and T. Jacks, “Restoration of p53 function leads to tumour regression in vivo,” Nature, vol. 445, no. 7128, p. 661–665, 2007.

[40] J. C. Acosta, A. Banito, T. Wuestefeld, A. Georgilis, P. Janich, J. P. Morton, D. Athineos, T.-W. Kang, F. Lasitschka, M. Andrulis, et al., “A complex secretory program orchestrated by the inflammasome controls paracrine senescence,” Nature cell biology, vol. 15, no. 8, p. 978–990, 2013.

[41] D. G. Burton and A. Stolzing, “Cellular senescence: immunosurveillance and future immunotherapy,” Ageing research reviews, vol. 43, p. 17–25, 2018.

[42] J. M. Valdez, L. Zhang, Q. Su, O. Dakhova, Y. Zhang, P. Shahi, D. M. Spencer, C. J. Creighton, M. M. Ittmann, and L. Xin, “Notch and tgfβ form a reciprocal positive regulatory loop that suppresses murine prostate basal stem/progenitor cell activity,” Cell stem cell, vol. 11, no. 5, p. 676–688, 2012.

[43] M. Hoare and M. Narita, “Notch and the 2 sasps of senescence,” Cell cycle, vol. 16, no. 3, p. 239–240, 2017.

[44] Y. V. Teo, N. Rattanavirotkul, N. Olova, A. Salzano, A. Quintanilla, N. Tarrats, C. Kiourtis, M. Müller, A. R. Green, P. D. Adams, et al., “Notch signaling mediates secondary senescence,” Cell Reports, vol. 27, no. 4, p. 997–1007, 2019.

[45] D. Paramos-de Carvalho, A. Jacinto, and L. Saúde, “The right time for senescence,” Elife, vol. 10, p. e72449, 2021.

